# Unveiling the benefits and gaps of wild pollinators on nutrition and income

**DOI:** 10.1101/2023.08.09.552641

**Authors:** Gabriela T. Duarte, Richard Schuster, Matthew G.E. Mitchell

## Abstract

Insect pollinators play a crucial role in global crop production, enhancing fruit and seed yield, improving fruit quality, and increasing crop nutritional value. However, wild pollinator populations worldwide have been experiencing alarming declines. This study investigates the contribution of wild pollinators to nutrition and farmers’ income in Canada, while examining the spatial distribution of these services. Using publicly available data on crop types, yields, nutrient content, and farm gate values, alongside information on natural habitats, we estimated the benefits of wild pollinators across national agricultural landscapes. Our findings reveal that these pollinators sustain approximately 24.4 million equivalent people each year in terms of nutrition and generate an annual income of nearly CAD$2.8 billion for farmers. However, significant benefit gaps exist due to the lack of nearby pollinator habitats and insufficient pollination of dependent crops on a national scale. Addressing these gaps could provide an additional nutrition supply for nearly 30 million equivalent people and increase farmer income by CAD$3 billion. We discuss how and where efforts focused on preserving and enhancing wild pollinator habitats, promoting sustainable farming practices, and raising awareness among stakeholders are crucial for the long-term viability of wild pollinator populations and the sustainability of agricultural systems. This research underscores the urgent need for a national strategy aimed at safeguarding wild pollinators. Implementing such a strategy would not only contribute to strengthening local economies in the regions highlighted in our work but also ensure the production of nutritionally essential food.

## Introduction

Insect pollinators, including bees, wasps, flies, and butterflies, play an essential role in crop production worldwide. Studies have shown that the presence of insect pollinators in crop fields can increase fruit and seed production, improve fruit quality, and enhance the nutritional value of crops (Klein *et al* 2007, IPBES 2016). For instance, blueberry and canola fields in Canada with high densities of bees have higher fruit/seed set and yields (Sabbahi *et al* 2005, Desjardins and Oliveira 2006, Button and Elle 2014). In fact, it has been estimated that more than three-quarters of the main global food crops rely to some extent on pollinator species (IPBES 2016). Hence, safeguarding their populations is crucial to maintain and improve agricultural production in addition to farmer livelihoods.

In some cases, native pollinators can be more effective at pollinating some crops than non-native species (Garibaldi *et al* 2013). For example, bumble bee species can “buzz” pollinate flowers of numerous crop species, significantly increasing fruit set and weight compared to European honeybees (Cooley and Vallejo-Marín 2021). Also, the presence of diverse communities of pollinator is more likely to generate yield stability and avoid production declines than less diverse communities, since pollinator species differ in their foraging behaviour, activity patterns and responses to changing conditions (Garibaldi *et al* 2011, 2014). However, both the richness and visitation rates of pollinator species are inversely related to the distance between crops and the natural or semi-natural ecosystems that provide nesting habitat (Ricketts *et al* 2008). Therefore, farmers should maintain and restore a variety of natural habitats near croplands to promote pollination services and increase the resilience of agricultural landscapes (Duarte *et al* 2018, 2020).

Despite their clear advantages for food production, wild pollinator populations have been declining worldwide, especially due to habitat loss, pesticide use, climate change, and pathogens (IPBES 2016). Global attention is now focused on the best way to halt this decline and guarantee their protection to help sustain many farmers’ livelihoods (Dicks *et al* 2016). As the level of dependency on animal pollination varies among crop types, regional agricultural economies will differ in their susceptibility to wild pollinator loss. For instance, roughly 20% of cultivated lands in Canada depend to some extent on animal pollinators and are concentrated in the central and southern parts of the country (Richards and Kevan 2002). However, because no national-scale analysis of pollination services has been completed for Canada, any pollination-friendly initiative or policy will lack the knowledge to target actions to most effectively maintain or improve pollination services.

Identifying sites where the contribution of wild pollinators is highest or that have the greatest potential to sustainably increase natural pollination and agricultural production is critical for prioritizing conservation and restoration efforts towards food security and biodiversity protection. Previous studies have combined biophysical and socio-economic data to assess pollination services at global or local scales (Chaplin-Kramer *et al* 2019, Koh *et al* 2016, Lautenbach *et al* 2012), while others have evaluated multiple ecosystem services at the national level but excluded pollination (Mitchell *et al* 2021). However, effective decision-making requires up-to-date and context-specific evidence-based information. Building on global models (Chaplin-Kramer *et al* 2019, 2023), our paper aims to understand how much wild pollinators in Canada potentially contribute to people in terms of nutritional values and farmers’ income and provide insight into the spatial distribution of these pollination benefits.

## Methods

We estimated the benefits of wild pollinators in agricultural landscapes across Canada at 240 m spatial resolution. To do this, we used publicly available data on crop type locations, yields, nutrient content, and farm gate values, as well as the location of natural habitats. We considered demand sites for wild pollinator services to be the croplands that rely on pollinator species to provide nutritional and economic benefits to people. Hence, service providing areas are the nearby natural areas that serve as habitat for these pollinator species. Our approach followed the methods of Chaplin-Kramer *et al* (2019), with modifications for Canadian landscapes, data, and to calculate economic benefits.

### Land use/cover data

We used land use and land cover (LULC) data from the Annual Crop Inventory (Agriculture and Agri-Food Canada 2022a), which uses optical and radar-based satellite imagery to produce national agricultural maps at 30m of spatial resolution. We chose to use the 2019 data because the complete sampling for more recent years (2020-2021) with ground data collection for specific crop types at certain locations in several provinces was not completed due to COVID-19 restrictions. By combining the LULC maps of each province into a mosaic, we obtained a single national-scale map at 30 m resolution. We resampled it to coarser resolution by calculating the proportion of each LULC class within each 240 m pixel. See more details in SM1 and Table S1.

### Crop pollination dependency

Klein et al. (2007) classified more than 120 crops according to their dependency on animal-mediated pollination, defined as the percent yield reduction that occurs with inadequate pollination. The authors provided 5 categories and we took the averages for the ranges of pollination dependence in each category (0.95 for ‘essential’, 0.65 for ‘great’, 0.25 for ‘modest’, 0.05 for ‘little’, 0 for “no-reduction”). With this information, we were able to define which crops depend on pollinators and to what extent (Table S2).

### Pollination sufficiency

Due to extensive conversion activities, some agricultural landscapes do not have sufficient natural habitat for a pollinator community that can adequately service nearby crops. We followed Chaplin-Kramer *et al*. (2019) and defined pollinator habitat as any natural LULC class - shrubland, wetland, peatland, grassland, and forest. We then calculated the proportion of natural habitat within ∼2 km (due to the LULC maps spatial resolution we used 1920m) for each pixel of cropland to determine its pollination sufficiency (Chaplin-Kramer *et al* 2019, 2023). Table S3 contains the results of a sensitivity analysis with other distance values. We assumed that cropland pixels with > 30% natural habitat in the surrounding landscape receive sufficient wild pollination for their pollinator-dependent (Kremen 2005). Pixel values between 0 and 30% of natural habitat were linearly interpolated between 0 and 1 for pollinator sufficiency. All pixels with >30% of natural habitat were assigned a value of 1.

### Crop production

Crop yield data (t ha^-1^) for each of the LULC classes in each province are from Statistics Canada surveys (Statistics Canada 2022a, 2022b, 2022c). We calculated a single value per province that represents the average yield for each given crop between 2015 and 2021. We calculated crop production by multiplying these values by the pixel area and the proportion of the corresponding LULC class in each pixel.

### Refuse fraction, nutrient content and dietary requirements

We used estimated refuse fractions for each crop (e.g., peels, seeds) from the Canadian Nutrient File database (Health Canada 2015) and reduced the crop production by this value - i.e., we multiplied crop production by 1 minus the refuse fraction. The same database provided information on macro- and micronutrient content for each crop. We took values for energy (KJ/100g), vitamin A (retinol activity equivalents (RAE) - mcg/100g), and folate (dietary folate equivalents (DFE) - mcg/100g). After calculating each nutrient content in 1 metric tonne of each crop, we multiply these contents by crop production in each pixel, resulting in layers of energy and nutrient production.

We used dietary daily requirements per capita for each micronutrient for different demographic populations (Health Canada 2022) and averaged the values within each age group (0-14, 15-64, 65+) and sex combination. The recommended nutrient intake is the average daily dietary intake sufficient to meet the nutrient needs of almost all healthy individuals in each life stage. Following Chaplin-Kramer et al. (2019), we estimated that 5% of the female population aged 15-64 are pregnant or breastfeeding. We then converted the daily values to annual requirements. For energy demand estimates, we used values from the work of Chaplin-Kramer *et al* (2019)(Table S4). Finally, using Canadian population data for 2019 (Statistics Canada 2019), we then applied these demographic breakdowns to the annual dietary requirements table to produce a national weighted average of total dietary need for energy and each nutrient.

### Pollinator-dependent crop production and equivalent people fed

We multiplied all factors to calculate wild pollinator-dependent nutrient production (WPN) for each crop in each pixel, following Equation 1:

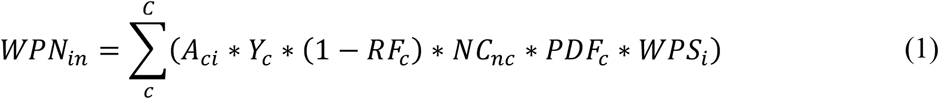

where A is the area (ha) occupied by the crop type “c” in pixel “ï”; Y is the potential yield of crop “c”, which was defined per Canadian province.; RF is the refuse fraction; NC is the nutrient content for nutrient “n” in crop “c”; PDF is the pollination dependency fraction for crop “c”; and WPS is the wild pollination sufficiency of pixel “ï”. We obtained the total wild pollinator-dependent nutrient production by summing the values across all crops for each pixel and for each nutrient.

These layers of nutritional benefits are expressed in different units of nutrient production (i.e., Kj, RAE, DFE). So, we modified these values by dividing each layer by the recommended annual dietary intake. Then, we averaged the resultant values in each pixel to obtain the equivalent number of people fed through pollination, which is the number of people whose nutritional needs could be fully met by wild pollinator-dependent crop production (Chaplin-Kramer *et al* 2019).

### Farm gate values

Farm gate value of crop production refers to the net value of a farm’s production as it leaves for the market, after selling costs have been subtracted. Using data from a variety of governmental sources (Statistics Canada 2022a, 2022b, 2022c), we estimated average farm gate values per metric tonne of each crop for each province between 2015-2021. Before averaging, all values were corrected for inflation using Consumer Prices Index, and 2019 as baseline (OECD 2022). We then calculated total farmer income related to wild pollinator-dependent crop production (WPV) using Equation 2:

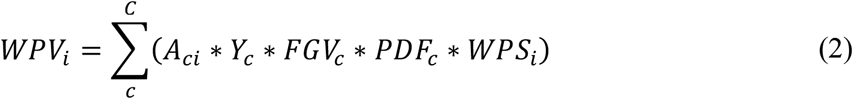

where FGV is the farm gate value per metric tonne of crop “c”.

### Mapping pollination benefits back to natural habitats

We mapped both the equivalent number of people fed and farmer income back to the surrounding pollinator habitat. We used a distance decay function (Equation 3), considering the distance between each focal crop pixel “j” and the surrounding natural habitat “i” to weight the contribution of each habitat pixel in creating the benefit (Chaplin-Kramer *et al* 2023). This weighted distance (Wdist_i_) considers only habitats within the same flying distance as the pollination sufficiency (dist_max_ of ∼2km) and was calculated following the habitat quality model of InVEST (Sharp *et al* 2020). We assumed that more distant habitats would have a lower weight because fewer wild pollinators from these habitats would reach the cultivated areas.

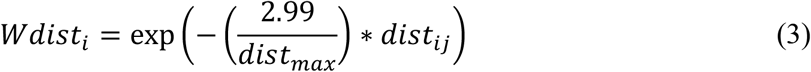

Along with distance, we factored in the area of natural habitat surrounding each focal crop pixel to distribute a portion of the pollination benefit back to these habitats. That is, we assumed that each natural habitat within the surrounding landscape contributes to a fraction of the benefit provision. This fraction is directly proportional to the habitat size and inversely proportional to its distance from the focal crop pixel. Therefore, we use Equation 4 to calculate the weighted benefit fraction for each focal pixel with cropland (Bweighted_j_). Finally, using Equation 5, we calculated the total benefit value of each pixel with natural land (Bnat), which is now the focal pixel, by considering all surrounding crop pixels and their weighted benefit fractions. Hence, we mapped the benefits back to the natural habitat accounting for the distance and area that it occupies.

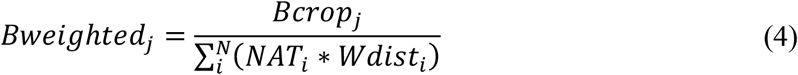

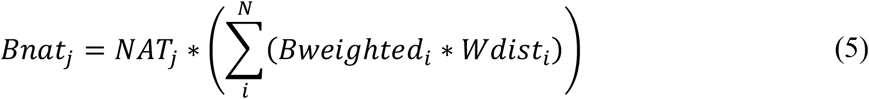

In these equations, “j” is the focal pixel with cropland in equation 4 and natural area in equation 5; “i” corresponds to the neighbor pixels with natural areas in equation 4 and croplands in equation 5; Bcrop corresponds to the benefit value of a pixel with cropland, derived from the wild pollinator-dependent crop production; NAT is the natural area in a neighboring pixel; Wdist is the weighted distance between the focal and neighbor pixels; Bweighted is the weighted benefit fraction; and Bnat would be the equivalent people fed value or farmers income (in CAD$) values of a natural area.

### Benefit gap

The term “benefit gap” refers to the portion of people’s needs not fulfilled by nature (Chaplin-Kramer *et al* 2019). In this case, it represents the potential loss of both equivalent number of people fed and farmer income due to insufficient wild pollination. We calculated the benefit gap for each pixel by subtracting the wild-pollination benefit values from the total potential benefit that would be possible with sufficient pollination, which is derived from the total pollinator-dependent crop production – that is, results of equations 1 and 2 without the WPS_i_ term.

### Data aggregation into Ecodistricts

We used ecodistricts as the subdivisions of Canadian agricultural areas to aggregate our findings and identify areas where different management actions for pollinators might be required. An ecodistrict is characterized by “distinctive assemblages of relief, landforms, geology, soil, vegetation, water bodies and fauna” (Agriculture and Agri-Food Canada 2013). The results presented here only consider ecodistricts that contain cropland or natural areas that benefit them (N = 417). Since each ecodistrict is different in size, for comparison purposes between them, we have divided the aggregated benefit values by the total service providing area. That is, the total natural area surrounding pollinator-dependent crops. Accordingly, we also divided the aggregated benefit gaps by the total pollinator-dependent crop area.

## Results

The estimated number of equivalent people fed each year in Canada that depends on wild pollinators is 24.4 million. Of this total, pollinator-dependent crop production in Saskatchewan feeds the most equivalent people (6.7 million), followed by Ontario (5.8 million) and Alberta (5.0 million). Additionally, almost CAD$2.8 billion is generated each year as income for farmers due to the presence of wild pollinators in Canada. Saskatchewan ($922M), Alberta ($693M) and Manitoba ($386M) are the provinces with the highest income value associated to wild pollinators and their natural habitats. On the other hand, lack of nearby pollinator habitat and pollination to dependent crops nationally leads to significant benefit gaps that if addressed could help generate an additional nearly 30 million equivalent people fed and CAD$ 3 billion in farmer income every year. Saskatchewan has the highest potential to increase benefits from pollinators for both people fed (11.5 million people) and income ($1.6B) benefit gaps. Table 1 summarizes these results per province and Figure S1 show their location.

**Table 1.** Benefit values and gaps per province.

| Province | Equivalent people fed<br>(thousand) | | Farmer's income<br>(CAD\$ million) | | Service<br>providing<br>areas<br>(million<br>hectares) | Pollinator-dependent crop<br>area (million<br>hectares) |
| --- | --- | --- | --- | --- | --- | --- |
|  | Benefit | Gap | Benefit | Gap |  |  |
| Newfoundland<br>and Labrador | 0.0 | 0.0 | 0.1 | 0.0 | 0.01 | $66 \times 10^{-6}$ |
| Prince Edward<br>Island | 110.5 | 12.6 | 12.1 | 0.7 | 0.28 | 0.03 |
| Nova Scotia | 36.5 | 8.0 | 23.4 | 2.0 | 0.86 | 0.02 |
| New<br>Brunswick | 35.7 | 0.6 | 22.2 | 0.0 | 0.83 | 0.02 |
| Quebec | 2,400.2 | 1,529.5 | 234.1 | 61.3 | 3.22 | 0.50 |
| Ontario | 5,838.0 | 7,090.4 | 249.2 | 298.4 | 2.84 | 1.44 |
| Manitoba | 4,060.7 | 5,473.3 | 385.6 | 476.8 | 3.10 | 2.35 |
| Saskatchewan | 6,689.9 | 11,490.9 | 922.3 | 1,574.0 | 6.95 | 7.18 |
| Alberta | 5,032.3 | 4,306.6 | 693.5 | 596.5 | 8.89 | 3.73 |
| British<br>Columbia | 178.9 | 67.5 | 244.2 | 79.3 | 1.08 | 0.11 |
| TOTAL | 24,382.8 | 29,979.3 | 2,786.7 | 3,089.0 | 28.07 | 15.37 |

**Table 2.** Benefit values per habitat type and their corresponding pollination service providing area.

| Habitat type | Equivalent people fed (thousand) | Farmer's income (CAD\$ million) | Service providing area (million hectares) |
| --- | --- | --- | --- |
| Shrubland | 2,013.3 | 235.5 | 2.39 |
| Wetland | 4,600.2 | 563.1 | 3.52 |
| Peatland | 5.2 | 0.8 | 0.01 |
| Grassland | 5,359.4 | 695.4 | 7.63 |
| Coniferous forest | 872.0 | 243.1 | 2.78 |
| Broadleaf forest | 9,060.5 | 827.7 | 7.92 |
| Mixedwood forest | 2,472.2 | 221.1 | 3.81 |

In 2019, Canada had 45.77 million hectares of agricultural land, encompassing crops, pastures, fallow areas, and related land uses. Our analysis focused on 34.86 million hectares of cropland, accounting for 76% of the total agricultural area. We examined data for 38 crop types, of which 17 rely on animal pollinators to some extent, covering an area of approximately 15.4 million hectares (Figure S2a). Canola/rapeseed, soybeans, and peas are the major pollinator-dependent crops, occupying 10, 2.6, and 1.9 million hectares respectively. The pollinator-dependent crops are distributed in 382 ecodistricts of the total 417 analysed, with 158 ecodistricts having more than half of their crop area relying on pollinators (Figure 1a).

**Figure 1.**
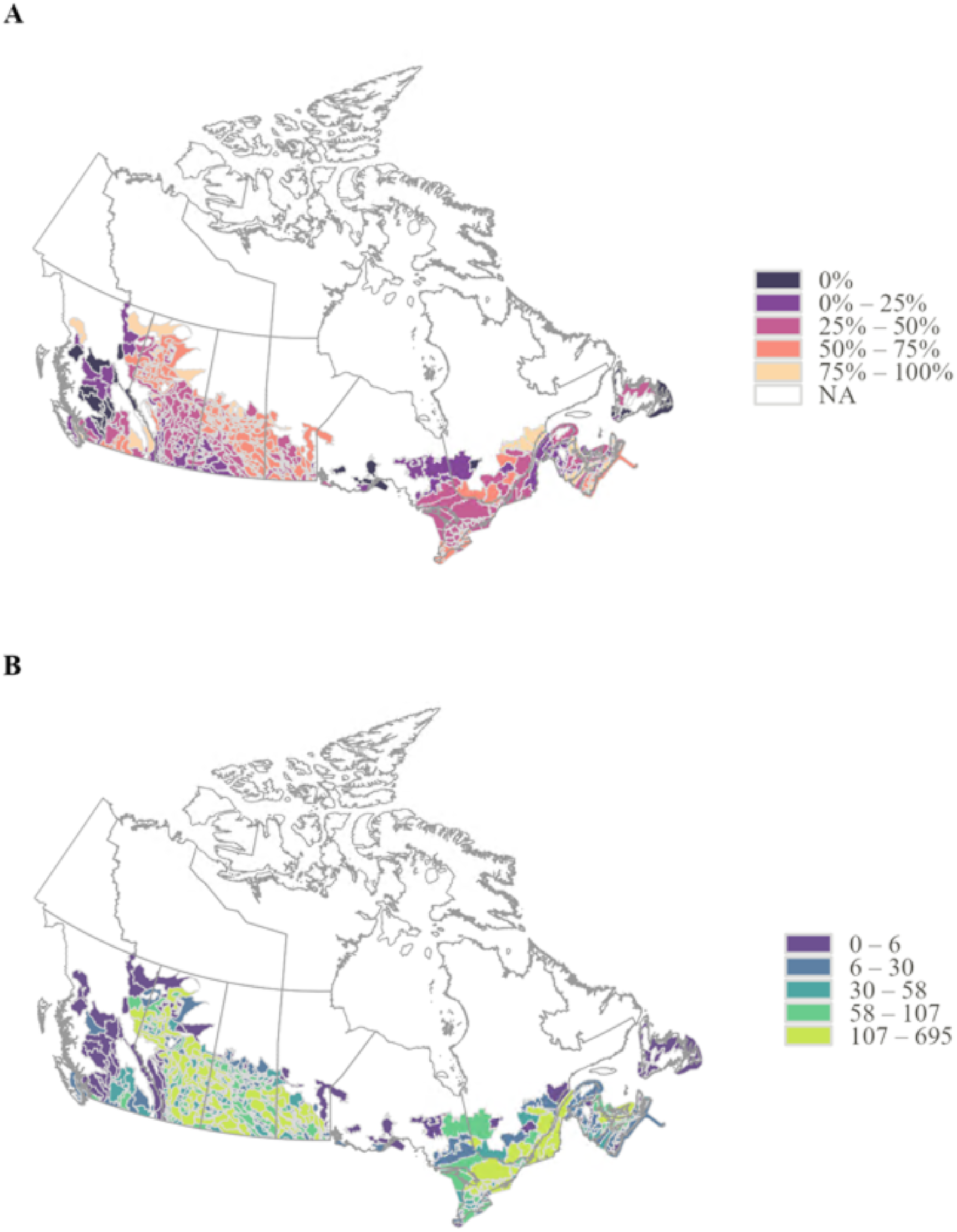
Crop dependency and service providing areas per ecodistrict. Maps showing: a) the percentage of the total crop area that is pollinator-dependent; and b) the total service providing area (thousand hectares). * NA values correspond to ecodistricts that have natural areas providing pollination services to neighbouring ecodistricts, even though they don’t contain crops themselves.

There are 84 ecodistricts that rank in the top 25% when it comes to the production per hectare of natural habitat for both equivalent people fed and farmer income benefits. The 6.6 million hectares of natural areas providing pollinator services within these 84 ecodistricts account for 60% of the national total number of equivalent people fed and 55% of the total farmer’s income. Thus, they are critical natural habitats that could be low-hanging fruits for conservation actions, as they provide, on average, high levels of pollination services in terms of nutrition and economic values. The locations of these ecodistricts are shown in light-bright-green in Figure 2a, which are concentrated in the central portion of the Prairies, southern Ontario, and the Saint Lawrence lowlands in Quebec.

**Figure 2.**
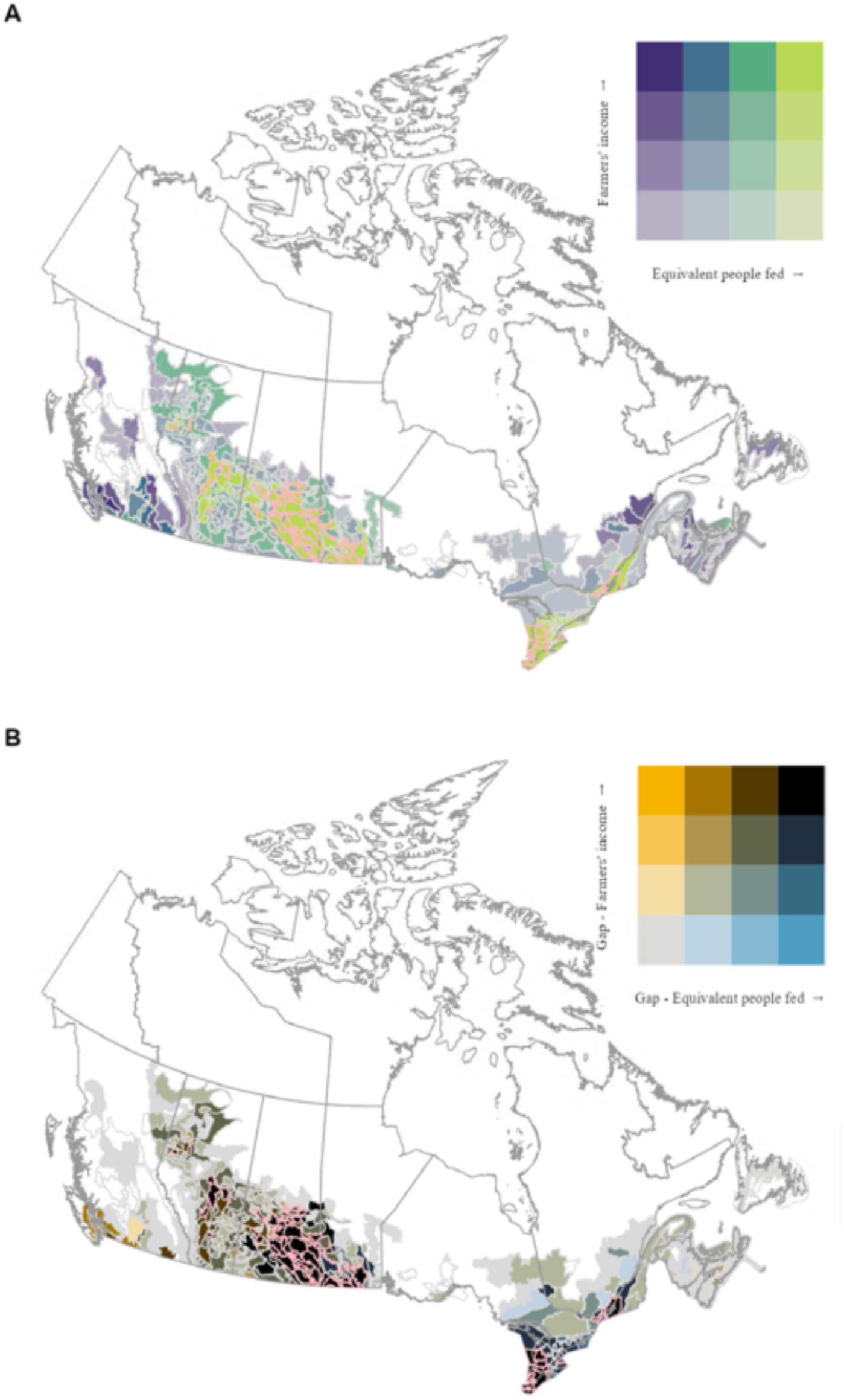
Benefits and gaps per ecodistrict. Maps showing the distribution of ecodistricts with different levels of a) benefit provision per service providing area; and b) benefit gap per area of pollinator-dependent crops. Ecodistricts with the highest levels in both maps (light-bright-green in a and black in b) have a light-pink outline.

We also identified possible targeted locations for restoration actions to increase pollinator habitat and benefits from pollination. They correspond to the 68 ecodistricts among the top 25% with the highest scores for both benefit gaps. These ecodistricts contain 7.6 million hectares of pollinator-dependent crops with insufficient wild pollination, which account for 70% of the national nutritional gap and 66% of the national economic gap. Their location is shown in black in Figure 2b. Additionally, there are 50 ecodistricts that are priority for both conservation and restoration actions to enhance pollinator services, and they are located mostly in Saskatchewan and southern Manitoba, represented with a light-pink outline in Figure 2a and b.

Our findings indicate that forests contribute to the largest share of both benefits among different habitat types. They help feed the equivalent of 12.4 million people and generate CAD$1.3 billion per year for farmers. Their service providing areas extend across 14.5 million hectares, primarily surrounding croplands in southern Ontario and Quebec, known for their high yield values, but also across the Prairie region (Figure 1b). Grasslands and wetlands also play crucial roles in providing pollination services in Canada. Together, they help feed nearly 10 million people and generate CAD$1.3 billion for farmers, with a significant impact in the Prairie region (Figure 1b).

## Discussion

Our study highlights the crucial role that wild pollinators play in food security and sustaining farmers’ income in Canada. To provide context, the estimated equivalent number of people with nutritional needs fully met by these pollinators exceeds half of Canada’s population (Statistics Canada 2019). Although, in reality, a significant portion of crop production is allocated for export, industrial use, or as feed grains, wild pollinators benefit a far greater number of people than estimated here by partially contributing to their nutritional needs, as an average person’s diet contains multiple sources of nutrients and energy. In addition, the estimated economic contributions from pollinators account for 8.5% of Canada-wide farmer revenues derived from the sale of principal field crops (including both pollinator-dependent and non-pollinator-dependent crops; Agriculture and Agri-Food Canada 2022b), and 5% of the total crop-related income (including more minor crops) estimated in our study. These findings and proportions align with the expected global decline in crop production in the absence of animal pollination, which is between 3 and 8% (Aizen *et al* 2009).

Our estimates demonstrate that Canada stands to gain significantly by promoting the protection of wild pollinators. Financial mechanisms and environmental programs that encourage cooperation among farmers could support this transition (Goldman and Tallis 2009, Ayambire *et al* 2021), as restoring natural habitats often benefits crop production for multiple landowners throughout the surrounding landscape and contributes to the establishment of a landscape structure capable of sustaining pollinator services (Duarte *et al* 2018, 2020). In our study, we identified specific locations where agri-environmental programs could prioritize investments by evaluating ecodistricts that offer high potential for promoting high levels of pollinator services through the conservation and/or restoration of natural habitats (ecodistricts in light-bright-green in Figure 2a and in black in Figure 2b). Within each of these ecodistricsts, the fine-scale results produced in this work (in 240m resolution) could be used to inform any participatory planning derived from these programs (Figure S2 and S3).

Investments in wild pollinator habitat also hold the potential to serve as climate solutions. Our findings highlight the critical role of Canadian forest ecosystems in providing pollination services, indicating a potential win-win scenario where conserving and restoring pollinator habitats can simultaneously enhance carbon storage in plants biomass (Drever *et al* 2021). This aspect could also further increase the eligibility of farmers for funding through the recently introduced federal Agricultural Climate Solutions program (Agriculture and Agri-Food Canada 2021). This and other similar funding sources could help compensate restoration implementation costs to create new natural habitats for pollinators. Additionally, it is important to note that other ecosystems can also provide significant co-benefits. For instance, our analysis emphasizes the crucial role that grasslands and wetlands also play in promoting pollination services. Unfortunately, these ecosystems have experienced considerable historical loss and degradation in Canada (Federal, Provincial and Territorial Governments of Canada 2010). Reversing this trend can provide substantial benefits for people, encompassing not the ones demonstrated in this study, but also carbon sequestration, water regulation and cultural values (Saleh and Karwacki 1996, Acreman and Holden 2013, Pattison-Williams *et al* 2018, Drever *et al* 2021).

A significant challenge lies in ensuring food security for an expanding global population while simultaneously preserving biodiversity and its contributions to human well-being (Chappell and LaValle 2011, Brussaard *et al* 2010). Our work underscores ecodistricts with high percentage of croplands that dependent on pollinators (Figure 1), and with high potential benefit provision (Figure 2). In these ecodistricts, going beyond habitat conservation and restoration is crucial to avoid the loss of current pollinator communities while increasing agriculture productivity. A promising approach consists in promoting ecological intensification in these croplands through management practices that conserve biodiversity beneficial to agricultural production, such as intercropping, crop rotations, and on-farm diversification (Kovács-Hostyánszki *et al* 2017). Previous studies have demonstrated that this shift in management can result in no net loss or even improved yields, monetary and nutritional values of the produced crops (Pywell *et al* 2015, Garibaldi *et al* 2016)

Accordingly, our analysis reveals significant benefit gaps, suggesting a substantial opportunity for a positive and profitable shift in agricultural management practices with respect to pollinators in Canada. Strikingly, restoring pollinator habitat and increasing pollinator benefits to pollinator-dependent crops have the potential to provide more benefits than are currently realized in Canada, especially in Ontario, Manitoba, and Saskatchewan. The latter province warrants special attention, as it has the highest benefits provision and the highest benefit gaps, due to its large extent of pollinator-dependent crops (Table 1) such as canola/rapeseed and peas. However, the best approaches for increasing pollinator habitat will likely also be region-specific across Canada, both due to the ecology and pollinator diversity, as well economic viability and physical constraints on-farm (Clearwater *et al* 2016). For example, in intensive agriculture areas like the Prairies, adding pollinator habitat might conflict with the use of large machinery. In high-value farmlands (e.g., southern Ontario) removing land from production to improve pollinator habitat could face financial constraints and require different incentives. Therefore, decision-makers should carefully evaluate the benefit gap estimates as their value could decrease if pollinator-dependent crops are restored to natural habitats.

Future refinements to our understanding of crop pollination across spatial scales are needed to better target conservation actions. By using a single flight distance and considering all natural areas as pollinator habitat, we simplify the diversity of pollinator species that occur in Canada (Agriculture and Agri-Food Canada 2014). This approach is necessary when conducting a nationwide modeling where species-specific data at this scale is largely lacking. We also did not consider the role of croplands as potential habitats for pollinator species, nor other local scale elements such as field margins and hedgerows. This could result in an overestimate of the value of natural habitat near croplands (Chaplin-Kramer *et al* 2019, 2023). Further fine scale analyses that build on our methods and results could modify flying distance and habitat preferences to correspond to the features of local and regional pollinator species (Sharp *et al* 2020). Last, there could be other unintended consequences when closing the benefit gaps that were not evaluated in this work, such as the appearance of pests due to the increased natural areas.

Our research highlights the pressing need to develop a national strategy aimed at safeguarding wild pollinators. Such a strategy has the potential to strengthen local economies and ensure the production of nutritionally essential food. Furthermore, our analysis can support the prioritization and implementation of conservation efforts aligned with Canada’s commitments under the recently agreed Kunming-Montreal Global Biodiversity Framework (Convention on Biological Diversity 2022). Goal B of this framework emphasizes the sustainable use and management of biodiversity, as well as the recognition, maintenance, and enhancement of nature’s contributions to people. By integrating practices that conserve pollinators with climate-smart agriculture and sustainable land management approaches, we can achieve several benefits, including enhanced biodiversity, improved crop production and resilience, and increased carbon storage within agricultural ecosystems.

## Acknowledgements

We are very grateful for the support and precious discussions provided by the NCC’s Conservation Policy and Planning team, and the M2L2 team. G.T.D. was founded by Nature Conservancy of Canada and M.G.E.M. was supported by an NSERC Discovery Grant.

