## Supplementary Information for "Unveiling the benefits and gaps of wild pollinators on nutrition and income"

**The PDF file includes:**

Figs. S1 to S3

Tables S1 to S4

Figure S1 – Map showing the location of the Canadian provinces and territories. The Prairies region corresponds to the provinces of Alberta, Saskatchewan and Manitoba.

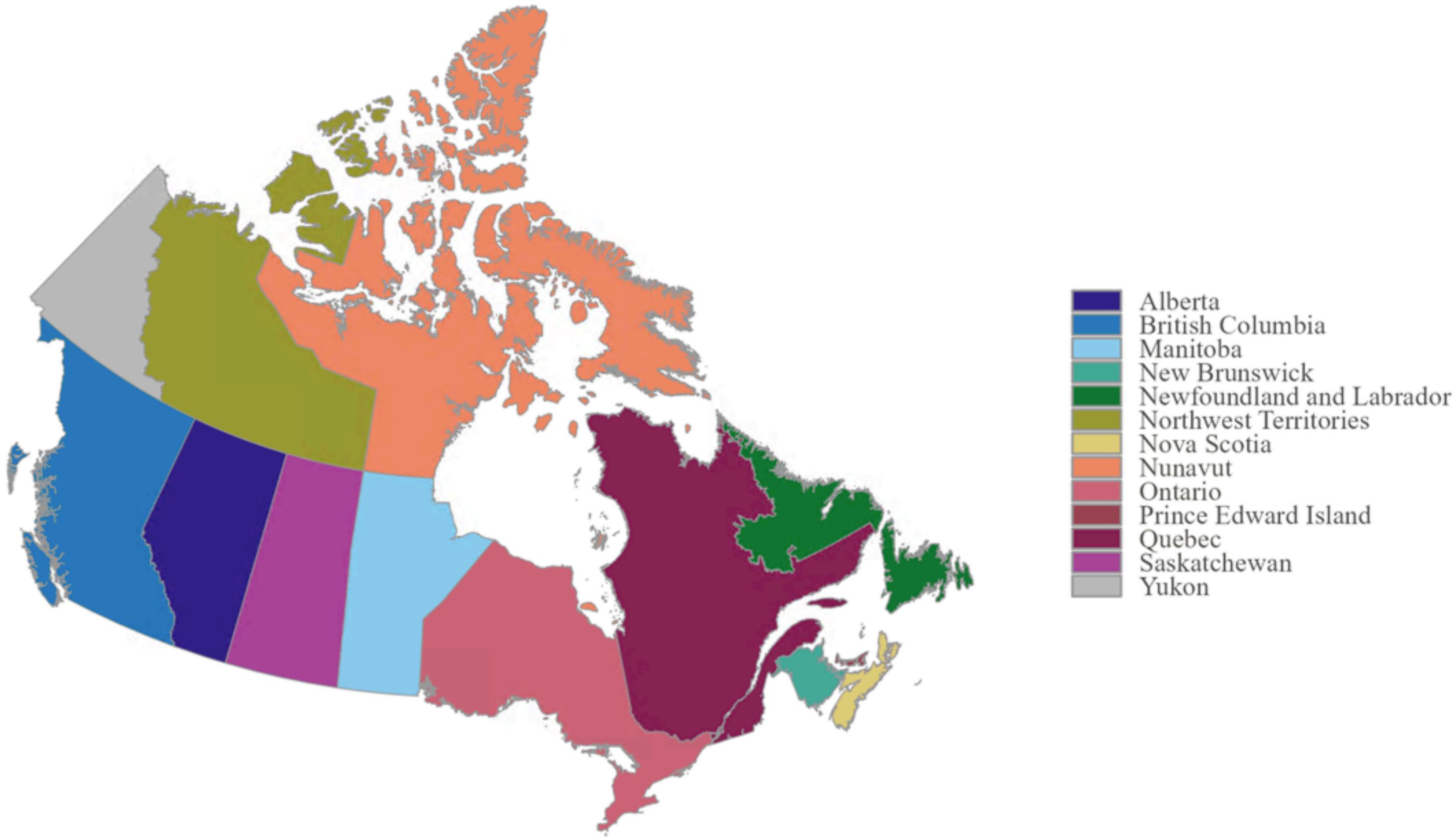

Figure S2 – Map showing the distribution of: a) pollinator-dependent crops (percentage of the pixel area) and b) natural areas within the pollinator flight distance (1920m) that serve as service provisioning areas (percentage of the pixel area).

**A**

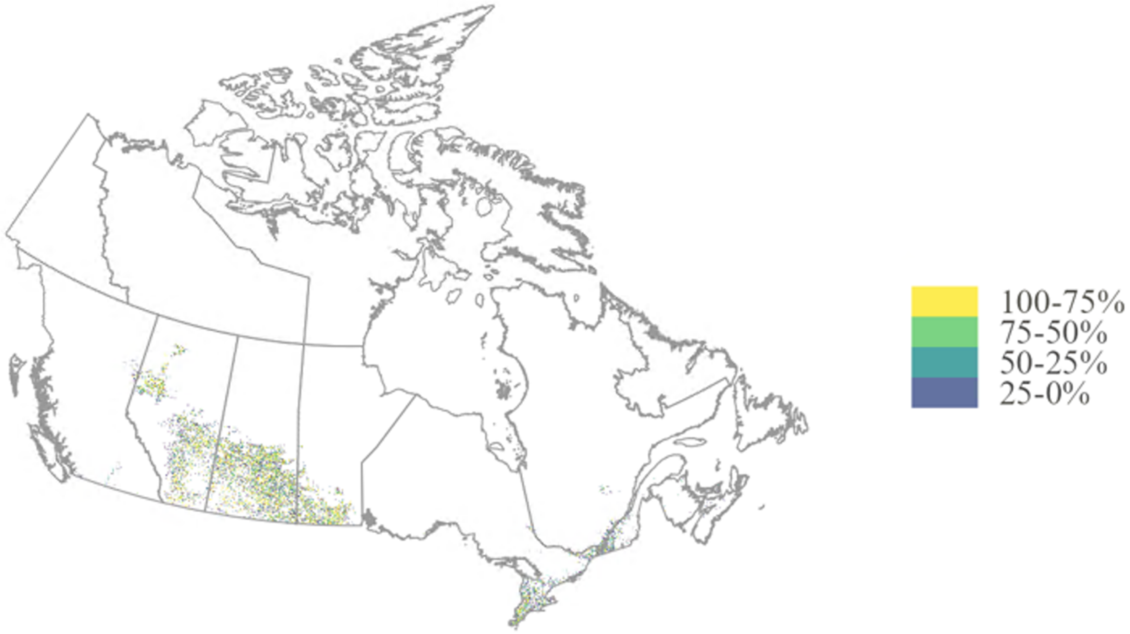


**B**

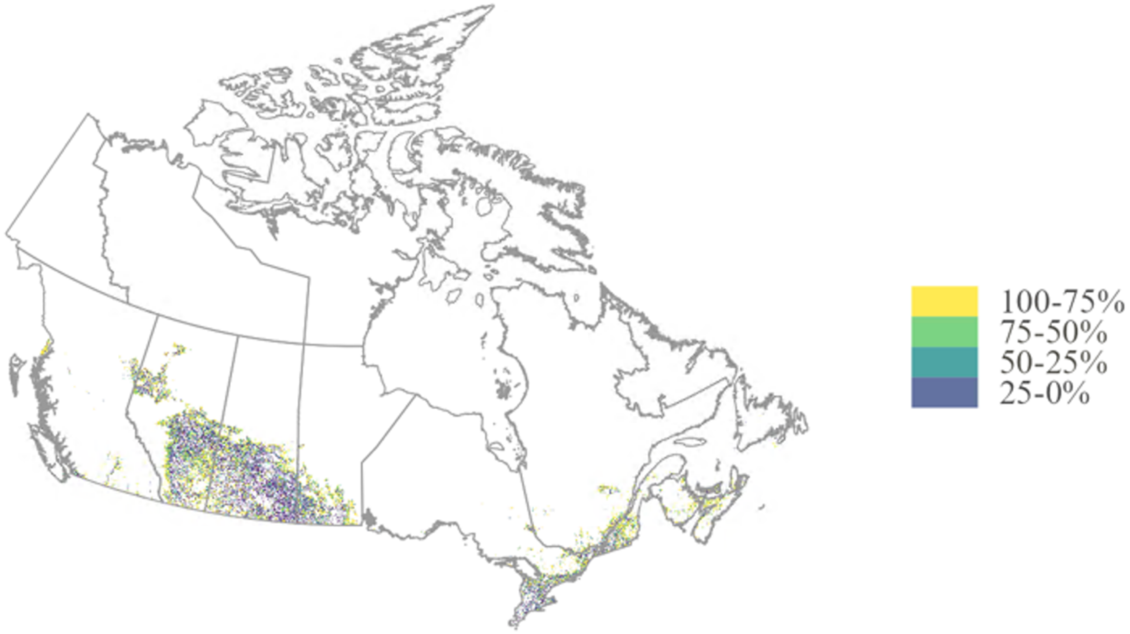

Figure S3 – Benefit and gap distribution. A and B provide, respectively, the equivalent number of people fed and farmer's income per pixel with natural areas. C and D provide their respectively benefit gaps per pixel with pollinator-dependent crop.

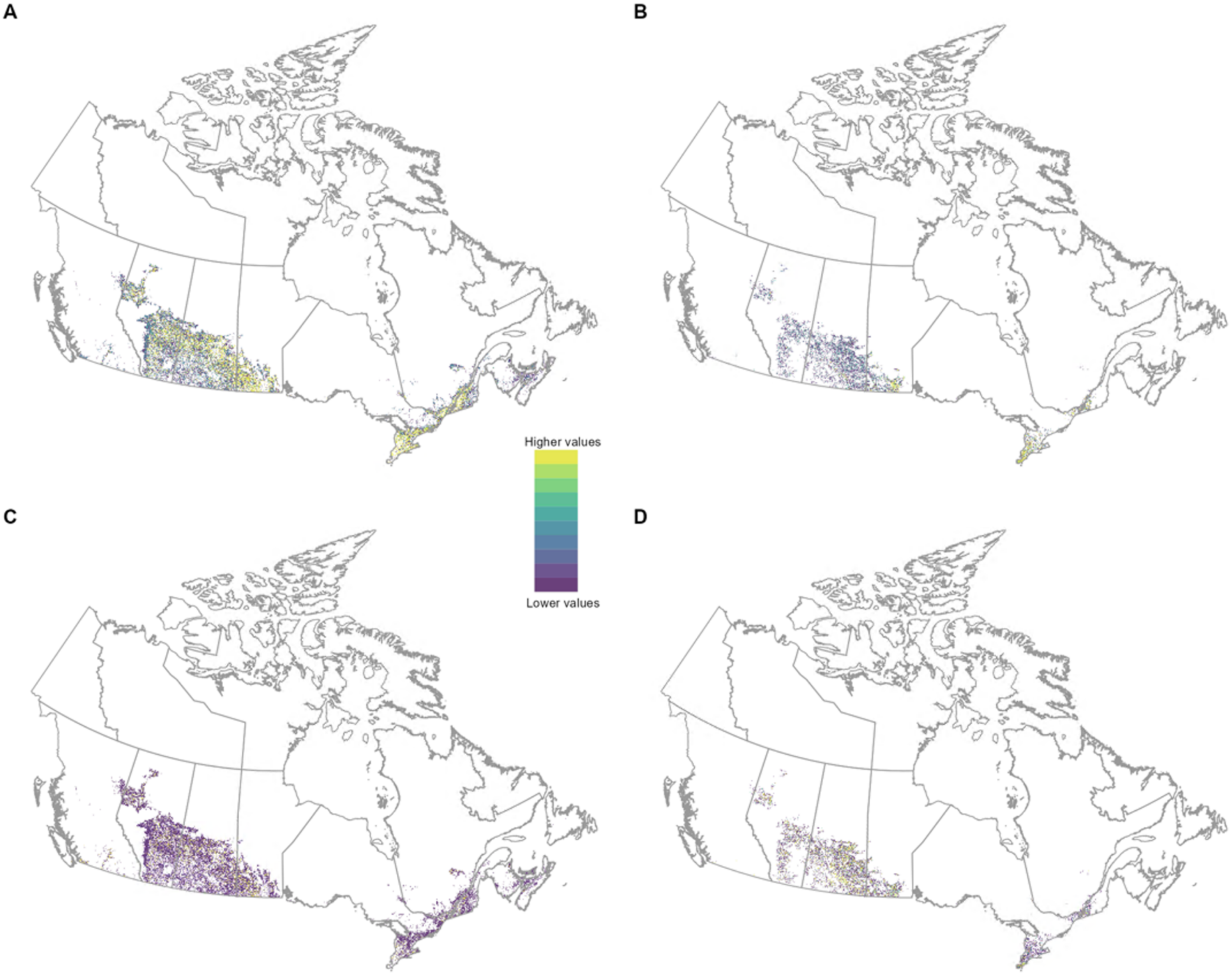

Transforming generic land use/land cover classes

The original land cover maps contained some generic crop group classes (e.g., "Cereals" and "Berries"). These classes were transformed into specific crop types based on which crops are primarily grown in each province for that group, measured by harvested area. When two or more crop types were considered as the main crops grown for a province, we averaged their yield, pollination dependence, nutrient content, and farm-gate values. However, these generic classes accounted for less than 0.15% of the total national crop area (Table S1).

Table S1 - Specific crop types used when transforming the generic land use classes, per province.

| **Provinces** | **Agriculture (undifferentiated)** | **Berries** | **Cereals** | **Other berry** | **Other crops** | **Other fruits** | **Other vegetables** |
|---|---|---|---|---|---|---|---|
| Newfoundland and Labrador | barley, millet, oats, rye, spelt, triticale, wheat, switchgrass, sorghum and quinoa | blueberries and cranberries | barley, millet, oats, rye, spelt, triticale, wheat, switchgrass, sorghum and quinoa | strawberries and saskatoon berries | barley, millet, oats, rye, spelt, triticale, wheat, switchgrass, sorghum and quinoa | blueberry, cranberry, apples and grapes | tomatoes, potatoes and sugarbeets |
| Prince Edward Island | barley, millet, oats, rye, spelt, triticale, wheat, switchgrass, sorghum and quinoa | blueberries and cranberries | barley, millet, oats, rye, spelt, triticale, wheat, switchgrass, sorghum and quinoa | strawberries and saskatoon berries | barley, millet, oats, rye, spelt, triticale, wheat, switchgrass, sorghum and quinoa | blueberry, cranberry, apples and grapes | tomatoes, potatoes and sugarbeets |
| Nova Scotia | barley, millet, oats, rye, spelt, triticale, wheat, switchgrass, sorghum and quinoa | blueberries and cranberries | barley, millet, oats, rye, spelt, triticale, wheat, switchgrass, sorghum and quinoa | strawberries and saskatoon berries | barley, millet, oats, rye, spelt, triticale, wheat, switchgrass, sorghum and quinoa | blueberry, cranberry, apples and grapes | tomatoes, potatoes and sugarbeets |
| New Brunswick | barley, millet, oats, rye, spelt, triticale, wheat, switchgrass, sorghum and quinoa | blueberries and cranberries | barley, millet, oats, rye, spelt, triticale, wheat, switchgrass, sorghum and quinoa | strawberries and saskatoon berries | barley, millet, oats, rye, spelt, triticale, wheat, switchgrass, sorghum and quinoa | blueberry, cranberry, apples and grapes | tomatoes, potatoes and sugarbeets |
| Quebec | barley, millet, oats, rye, spelt, triticale, wheat, switchgrass, | blueberries and cranberries | barley, millet, oats, rye, spelt, triticale, wheat, switchgrass, | strawberries and saskatoon berries | barley, millet, oats, rye, spelt, triticale, wheat, switchgrass, | blueberry, cranberry, apples | tomatoes, potatoes and sugarbeets |

| | | | | | | | |
|---|---|---|---|---|---|---|---|
| | sorghum and quinoa | | sorghum and quinoa | | sorghum and quinoa | and grapes | |
| Ontario | barley, millet, oats, rye, spelt, triticale, wheat, switchgrass, sorghum and quinoa | strawberries and blueberries | barley, millet, oats, rye, spelt, triticale, wheat, switchgrass, sorghum and quinoa | strawberries and saskatoon berries | barley, millet, oats, rye, spelt, triticale, wheat, switchgrass, sorghum and quinoa | blueberry, cranberry, apples and grapes | tomatoes, potatoes and sugarbeets |
| Manitoba | barley, millet, oats, rye, spelt, triticale, wheat, switchgrass, sorghum and quinoa | blueberries and cranberries | barley, millet, oats, rye, spelt, triticale, wheat, switchgrass, sorghum and quinoa | strawberries and saskatoon berries | barley, millet, oats, rye, spelt, triticale, wheat, switchgrass, sorghum and quinoa | blueberry, cranberry, apples and grapes | tomatoes, potatoes and sugarbeets |
| Saskatchewan | barley, millet, oats, rye, spelt, triticale, wheat, switchgrass, sorghum and quinoa | strawberries and saskatoon berries | barley, millet, oats, rye, spelt, triticale, wheat, switchgrass, sorghum and quinoa | strawberries and saskatoon berries | barley, millet, oats, rye, spelt, triticale, wheat, switchgrass, sorghum and quinoa | blueberry, cranberry, apples and grapes | tomatoes, potatoes and sugarbeets |
| Alberta | barley, millet, oats, rye, spelt, triticale, wheat, switchgrass, sorghum and quinoa | strawberries and saskatoon berries | barley, millet, oats, rye, spelt, triticale, wheat, switchgrass, sorghum and quinoa | strawberries and saskatoon berries | barley, millet, oats, rye, spelt, triticale, wheat, switchgrass, sorghum and quinoa | blueberry, cranberry, apples and grapes | tomatoes, potatoes and sugarbeets |
| British Columbia | barley, millet, oats, rye, spelt, triticale, wheat, switchgrass, sorghum and quinoa | blueberries and cranberries | barley, millet, oats, rye, spelt, triticale, wheat, switchgrass, sorghum and quinoa | strawberries and saskatoon berries | barley, millet, oats, rye, spelt, triticale, wheat, switchgrass, sorghum and quinoa | blueberry, cranberry, apples and grapes | tomatoes, potatoes and sugarbeets |

Table S2 – Pollination dependency values used per crop per province. NL = Newfoundland and Labrador, PE = Prince Edward Island, NS = Nova.Scotia, NB = New Brunswick, QC = Quebec, ON = Ontario, MB = Manitoba, SK = Saskatchewan, AB = Alberta, BC = British Columbia.

| **Land use class** | **NL** | **PE** | **NS** | **NB** | **QC** | **ON** | **MB** | **SK** | **AB** | **BC** |
|---|---|---|---|---|---|---|---|---|---|---|
| Agriculture (undifferentiated) | 0 | 0 | 0 | 0 | 0 | 0 | 0 | 0 | 0 | 0 |
| Barley | 0 | 0 | 0 | 0 | 0 | 0 | 0 | 0 | 0 | 0 |
| Beans | 0.1 | 0.1 | 0.1 | 0.1 | 0.1 | 0.1 | 0.1 | 0.1 | 0.1 | 0.1 |
| Berries | 0.65 | 0.65 | 0.65 | 0.65 | 0.65 | 0.4 | 0.65 | 0.25 | 0.25 | 0.65 |
| Blueberry | 0.65 | 0.65 | 0.65 | 0.65 | 0.65 | 0.65 | 0.65 | 0.65 | 0.65 | 0.65 |
| Borage | 0.65 | 0.65 | 0.65 | 0.65 | 0.65 | 0.65 | 0.65 | 0.65 | 0.65 | 0.65 |
| Buckwheat | 0.65 | 0.65 | 0.65 | 0.65 | 0.65 | 0.65 | 0.65 | 0.65 | 0.65 | 0.65 |
| Camelina | 0.05 | 0.05 | 0.05 | 0.05 | 0.05 | 0.05 | 0.05 | 0.05 | 0.05 | 0.05 |
| Canaryseed | 0 | 0 | 0 | 0 | 0 | 0 | 0 | 0 | 0 | 0 |
| Canola/rapeseed | 0.4 | 0.4 | 0.4 | 0.4 | 0.4 | 0.4 | 0.4 | 0.4 | 0.4 | 0.4 |
| Cereals | 0 | 0 | 0 | 0 | 0 | 0 | 0 | 0 | 0 | 0 |
| Chickpeas | 0 | 0 | 0 | 0 | 0 | 0 | 0 | 0 | 0 | 0 |
| Corn | 0 | 0 | 0 | 0 | 0 | 0 | 0 | 0 | 0 | 0 |
| Cranberry | 0.65 | 0.65 | 0.65 | 0.65 | 0.65 | 0.65 | 0.65 | 0.65 | 0.65 | 0.65 |
| Fababeans | 0.25 | 0.25 | 0.25 | 0.25 | 0.25 | 0.25 | 0.25 | 0.25 | 0.25 | 0.25 |
| Flaxseed | 0.05 | 0.05 | 0.05 | 0.05 | 0.05 | 0.05 | 0.05 | 0.05 | 0.05 | 0.05 |
| Hemp | 0 | 0 | 0 | 0 | 0 | 0 | 0 | 0 | 0 | 0 |
| Lentils | 0 | 0 | 0 | 0 | 0 | 0 | 0 | 0 | 0 | 0 |
| Millet | 0 | 0 | 0 | 0 | 0 | 0 | 0 | 0 | 0 | 0 |
| Mustard | 0.25 | 0.25 | 0.25 | 0.25 | 0.25 | 0.25 | 0.25 | 0.25 | 0.25 | 0.25 |
| Oats | 0 | 0 | 0 | 0 | 0 | 0 | 0 | 0 | 0 | 0 |
| Orchards | 0.65 | 0.65 | 0.65 | 0.65 | 0.65 | 0.65 | 0.65 | 0.65 | 0.65 | 0.65 |
| Other berry | 0.25 | 0.25 | 0.25 | 0.25 | 0.25 | 0.25 | 0.25 | 0.25 | 0.25 | 0.25 |
| Other crops | 0 | 0 | 0 | 0 | 0 | 0 | 0 | 0 | 0 | 0 |
| Other fruits | 0.4 | 0.4 | 0.4 | 0.4 | 0.4 | 0.4 | 0.4 | 0.4 | 0.4 | 0.4 |
| Other vegetables | 0 | 0 | 0 | 0 | 0 | 0 | 0 | 0 | 0 | 0 |
| Peas | 0.025 | 0.025 | 0.025 | 0.025 | 0.025 | 0.025 | 0.025 | 0.025 | 0.025 | 0.025 |

| | | | | | | | | | | |
|---|---|---|---|---|---|---|---|---|---|---|
| Potatoes | 0 | 0 | 0 | 0 | 0 | 0 | 0 | 0 | 0 | 0 |
| Quinoa | 0 | 0 | 0 | 0 | 0 | 0 | 0 | 0 | 0 | 0 |
| Rye | 0 | 0 | 0 | 0 | 0 | 0 | 0 | 0 | 0 | 0 |
| Sorghum | 0 | 0 | 0 | 0 | 0 | 0 | 0 | 0 | 0 | 0 |
| Soybeans | 0.25 | 0.25 | 0.25 | 0.25 | 0.25 | 0.25 | 0.25 | 0.25 | 0.25 | 0.25 |
| Spelt | 0 | 0 | 0 | 0 | 0 | 0 | 0 | 0 | 0 | 0 |
| Spring wheat | 0 | 0 | 0 | 0 | 0 | 0 | 0 | 0 | 0 | 0 |
| Sugarbeets | 0 | 0 | 0 | 0 | 0 | 0 | 0 | 0 | 0 | 0 |
| Sunflower | 0.25 | 0.25 | 0.25 | 0.25 | 0.25 | 0.25 | 0.25 | 0.25 | 0.25 | 0.25 |
| Switchgrass | 0 | 0 | 0 | 0 | 0 | 0 | 0 | 0 | 0 | 0 |
| Tomatoes | 0.05 | 0.05 | 0.05 | 0.05 | 0.05 | 0.05 | 0.05 | 0.05 | 0.05 | 0.05 |
| Triticale | 0 | 0 | 0 | 0 | 0 | 0 | 0 | 0 | 0 | 0 |
| Vineyards | 0 | 0 | 0 | 0 | 0 | 0 | 0 | 0 | 0 | 0 |
| Winter wheat | 0 | 0 | 0 | 0 | 0 | 0 | 0 | 0 | 0 | 0 |

Table S3 – Results of sensitivity analysis related to the national estimates for each benefit with using different flight distances. The percentage of change is demonstrated in comparison to the chosen baseline distance.

| **Flight distance (m)** | **Percentage of change (%)** | | |
|---|---|---|---|
| | **Distance** | Equivalent people fed | **Farmer's income** |
| 960 | -50 | -16 | -15 |
| 1200 | -38 | -10 | -9 |
| 1440 | -25 | -6 | -6 |
| 1680 | -12 | -3 | -3 |
| 1920 | baseline | baseline | baseline |
| 2160 | 12 | 3 | 3 |
| 2400 | 25 | 5 | 5 |
| 2640 | 38 | 7 | 6 |
| 2880 | 50 | 8 | 8 |

Table S4 – Annual nutrient requirements and total Canadian population (2019) per demographic group. Values correspond to energy (KJ), vitamin A (retinol activity equivalents (RAE)), and folate (dietary folate equivalents (DFE)).

| Age | Sex | RAE | DFE | KJ | Population |
|---|---|---|---|---|---|
| 0-14 | F | 182625 | 54178.74999 | 2339683.386 | 2941435 |
| 0-14 | M | 182625 | 54178.74999 | 2518483.488 | 3072854 |
| 15-64 | F | 261439.1929 | 148844.8538 | 3290227.518 | 12406989 |
| 15-64 | M | 328725 | 146100 | 4087951.05 | 12575373 |
| 65+ | F | 255675 | 146100 | 2866914.456 | 3562753 |
| 65+ | M | 328725 | 146100 | 3542381.508 | 3029858 |
